# New drinking water genome catalog identifies a globally distributed bacterial genus adapted to disinfected drinking water systems

**DOI:** 10.1101/2024.05.22.595307

**Authors:** Ashwin S Sudarshan, Zihan Dai, Marco Gabrielli, Solize Oosthuizen-Vosloo, Konstantinos T. Konstantinidis, Ameet J Pinto

## Abstract

Genome-resolved insights into the structure and function of the drinking water microbiome can advance the effective management of drinking water quality. To enable this, we constructed and curated thousands of metagenome-assembled and isolate genomes from drinking water distribution systems globally to develop a Drinking Water Genome Catalog (DWGC). The current DWGC disproportionately represents disinfected drinking water systems due to a paucity of metagenomes from non-disinfected systems. Using the DWGC, we identify core genera of the drinking water microbiome including a genus (UBA4765) within the order *Rhizobiales* that is frequently detected and highly abundant in disinfected drinking water systems. We demonstrate that this genus has been widely detected but incorrectly classified in previous amplicon sequencing-based investigations of the drinking water microbiome. Further, we show that a single genome variant (genomevar) within this genus is detected in >82% of drinking water systems included in this study. We propose a provisional name for this uncultured bacterium as “*Raskinella chlorumaquaticus*” and describe the genus as “*Raskinella*” (pending SeqCode approval). Metabolic annotation and modeling-based predictions indicate that this bacterium is capable of necrotrophic growth, able to metabolize halogenated compounds, proliferates in a biofilm-based environment and shows clear indications of disinfection-mediated selection.

**Synopsis:** Creation and analysis of a new curated drinking water genome catalog identifies uncharacterized bacterial species that is widely distributed in disinfected drinking water systems.

## Introduction

The drinking water microbiome^1^ is a diverse collection of bacteria and archaea^2^, eukaryotes^3^, and viruses^4^ and varies in composition spatially and temporally^5,6^ from source to tap^7^. Considering the myriad ways in which biological activity in drinking water infrastructure – from treatment to distribution and in the built environment – can affect the safety and aesthetic quality of drinking water^8,9^, understanding the structure and function of the drinking water microbiome is critical. One approach to develop generalizable insights involves direct comparative analysis of microbial community composition and associated selective pressures (e.g., disinfection, nutrient availability) across different systems (i.e., meta-analysis). While differences in methodological choices (e.g., DNA extraction or sequencing protocol) across studies can limit the quantitative utility of such analysis^10^, it is still possible to obtain important generalizable insights from such cross-study comparisons^1^.

Previous meta-analyses of the drinking water microbiome have relied on 16S rRNA gene amplicon sequencing datasets^10,11^. Both studies showed that the drinking water microbiome consists of a small core community that exhibits signs of selection from both water treatment and distribution practices. For instance, Thom et al. determined that the assembly of drinking water microbial communities post-disinfection was primarily deterministic^11^. This deterministic community assembly results in a community composition that can impact disinfectant residuals (e.g., via nitrification) and harbor bacterial genera of concern (i.e., *Legionella* and *Mycobacterium*). While amplicon sequencing studies have indeed provided extensive insights into the composition and biogeography of the drinking water microbiome, these results are inherently limited as they do not provide functional information. Functional information can shed light on processes governing phenomena like biocorrosion, biofilm formation, and interactions between microbial community members that could impact pathogen prevalence^12^ and also provide clues on how treatment and distribution practices exert selective pressures. Santos et al. (2016) highlighted that a significant portion of 16S rRNA gene sequences from the drinking water systems do not have genome representatives in reference databases and 16S rRNA gene sequence-based functional predictions are reliant on adequate genomic representation in reference databases; this makes such analysis questionable for drinking water studies^10^.

Metagenomic studies have identified specific metabolic and stress tolerance traits that may enable survival and growth in DWDSs. For instance, studies have shown enrichment of functional traits (e.g., necrotrophy) in a treatment and disinfection strategy-specific manner^13–16^. Metagenomics has also helped identify pathogenic traits^15^ and antibiotic resistant genes^17,18^ and further elucidated the prevalence of phages^4^ and anti-phage defense mechanisms^19^ which can impact microbial community dynamics. Recently, Liu et al. (2024)^15^ conducted a meta-analysis involving reconstruction and consolidation of metagenome assembled genomes (MAGs) from multiple drinking water metagenomes. They identified core organisms across studies, but their analysis was largely focused on (1) potential pathogenic bacteria and their likely associations with other community members and (2) on the metabolic traits of specific populations (e.g., comammox bacteria); they do not delve into the association between selective pressures shaping the drinking water microbiome and the functional potential of populations being selected.

The present study utilizes publicly available metagenomes to reconstruct and consolidate MAGs and develop an open-source Drinking Water Genome Catalog (DWGC) with the goal of identifying populations that are under selection in drinking water systems and its implications broadly on the microbial ecosystems in DWDSs. Through this analysis, this study finds that the core drinking water microbiome is highly structurally constrained. In the course of delineating the core drinking water microbiome, this study identified a globally distributed and potentially consequential, but consistently misannotated, bacterial genus and a highly abundant bacterial genomevar within it that is present in disinfected DWDS globally. The MAGs from this group were used to infer its ecology and model their potential metabolism to predict its niche within the DWDS as well as potential for regrowth and survival within this ecosystem.

## Materials and Methods

### Data retrieval and curation

Metagenomes were retrieved from studies indexed on Web of Science on or before September 5, 2022 using search string “drinking water” (All Fields) AND metagenom* (All Fields). Only metagenomes, MAGs or isolates genomes from finished water at the drinking water treatment plants (DWTP) or drinking water distribution system (DWDS) or point-of-use (PoU) were included in this study. This resulted in retrieval of raw reads for 208 metagenomes from 85 DWDSs (**Supplemental Table S1A**) and 55 isolate genomes from NCBI (**Supplemental Table S1B**). Metagenomes were grouped by DWDS with the exception of the Ke et al. (2022)^5^ datasets which were processed according to the sampling site by using one replicate per timepoint due to high redundancy between replicate metagenomes. MAGs were also generated from unpublished metagenomes from a Boston (USA) drinking water system^20^.

### Metagenome data processing and co-assembly

Adapters and poor-quality sequences were trimmed and filtered with fastp v0.20.1/v0.22.0/v0.23.2^21^ using the flags --qualified_quality_phred 20, --trim_poly_g, -- trim_poly_x and --length_required 20. Vector contamination in metagenomes was identified by mapping reads to the UniVec Core 10.0 database using BWA-MEM v0.7.17/BWA-MEM2 v2.2.1^22,23^, filtered using SAMtools v1.9/v1.16.1^24^ and reads were extracted using bedtools v2.30.0/samtools v1.16.1^24,25^. Subsequently, metagenomic reads from multiple locations within a single DWDS were combined for co-assembly. Metagenomic assembly was performed with MetaSPades v3.10.1/3.15.3/3.15.5 using a set of custom kmers (21,33,55,77)^26^. This resulted in a total of 85 metagenomic assemblies representing 85 DWDSs.

### Binning and refinement of MAGs

Quality filtered reads were mapped to contigs greater than 499 bps in their respective assemblies using BWA-MEM v0.7.17^22^ and bam files were sorted and indexed using SAMtools v1.3.1/v1.9/v1.16.1^24^. Contig coverage depth profiles for MetaBAT2 and VAMB were obtained using jgi_summarize_bam_contig_depths from bowtie2 v2.1.0/ MetaBAT2 v2.15^27,28^. Contigs were binned using CONCOCT^29^ for co-assemblies or MetaBAT2^28^ for single sample assemblies and VAMB^30^. CONCOCT binning was performed within Anvi’o v5.5/7.1^31^ using contigs larger than 2500 bps. MetaBAT2 v2.12.1/v2.15 binning was performed using contigs greater than 2500 bps and VAMB v3.0.8 was used with a minimum contig length of 2500 bps and minimum bin size of 200,000 bps. In the event that dereplicated MAGs from multiple binning approaches were available from a study^13,32,16^, we only performed VAMB based-binning prior to further dereplication. In some instances^20,33^, MAGs were used directly as they were generated using the workflow adopted in this study. The quality of the bins was estimated using CheckM v1.0.13/v1.2.2^34^ and bins with completeness > 50% and redundancy greater than 10% manually refined using anvi-refine from Anvi’o v5.5/7.1.

### Dereplication of bins and construction of the Drinking Water Genome Catalog (DWGC)

CheckM2 v0.1.3^35^ was used to evaluate the quality of bins prior to dereplication as it does not rely exclusively on marker genes to assess quality (see results and discussion section for further details). Bin quality was determined using the following formula: Quality Score = Completeness – 5 x Contamination. Bins with a minimum quality score of 50 were retained for further dereplication using dRep v3.4.0^36^. First, dereplication was performed with secondary alignment criterion of 0.99, minimum completeness of 50% and maximum contamination of 10% (-sa 0.99 - comp 50 -con 10) with FastANI^37^. This resulted in a non-redundant set of MAGs that constitute the DWGC. These non-redundant MAGs were further dereplicated using a secondary alignment criterion of 0.95 and coverage threshold of 0.3 to obtain representative MAGs at the species-level. Species-level representative MAGs were selected by calculating the quality score of each MAG within a species cluster using the following formula: Completeness – 5 x Contamination + 0.5 log(N50) + (centrality - S_ani) and the MAG with the highest score within a species cluster was selected as the representative MAG for that species cluster. This formula is a modification from Almeida et al. (2021)^38^ and emphasizes centrality weight to identify the most representative MAG within a given species cluster.

### Annotation and phylogenetic placement of MAGs

Taxonomic annotation of MAGs was performed using the classify workflow (classify_wf) from gtdb-tk v2.1.1 using the GTDB reference database release 207_v2^39^. Bacterial MAGs from the dataset were functionally annotated using Bakta v1.7^40^ using the flags –meta –compliant –keep-contig-headers and the archaeal MAGs from the dataset were annotated using Prokka v1.14.6^41^ using the flags –kingdom Archaea –metagenome –compliant to identify tRNAs, 5S, 16S and 23S rRNA genes from the genomes. MAGs were categorized as “High-Quality draft” and “Medium-Quality draft” according to the MIMAG/MISAG criteria^42^. High Quality MAGs were defined as those with a completeness > 90%, contamination < 5%, tRNAs for 18 out of the 20 amino acids, and the presence of 23S, 16S and 5S rRNA genes. MAGs that failed to satisfy the high-quality MAGs criteria but with completeness above 50% and contamination <10% are considered medium-quality MAGs. Multiple sequence alignment file from gtdb-tk^39^ was used to construct the phylogenetic tree for all bacterial MAGs. The alignment was trimmed using trimal v1.4.rev15^43^ using the flag -gappyout. The trimmed alignment file was used to construct a maximum likelihood phylogenetic tree with RAxML v8.2.12^44^ using the command raxmlHPC-PTHREADS with the PROTGAMMAWAG model and using seed 3301. Tree visualization and annotations were performed on iTOL v6^45^. Faith’s phylogenetic diversity^46^ for select groups within the DWGC was calculated using the constructed tree with the abdiv package^47^ on R^48^.

### Identification of core drinking water microbiome

Reads from all metagenomes were competitively mapped against all species-level representative MAGs using BWA-MEM v0.7.17^22^. Sorting and indexing of BAM files was performed using anvi-init-bam from Anvio v7.1^31^. CoverM v0.6.1^49^ was used with the following parameters: --min-read-percent-identity 0.95 --min-read-aligned-percent 0.75 and -m covered_fraction and relative_abundance to determine the relative abundance and covered fraction for each MAG in each metagenome; the latter parameter captures the proportional basepairs within a MAG with atleast one mapped read from the metagenome. By default, CoverM requires that at least 10% covered fraction for a MAG to be detected in a metagenome. The average relative abundance of a genus was calculated by dividing the cumulative relative abundance of all MAGs within that genus by the number of metagenomic assemblies in which that MAGs from that genus was detected. Further, we used a detection frequency threshold of 30% and 60% to evaluate the distribution of genera across metagenomic assemblies to identify core microbial taxa. Ecologically-relevant core taxa were also identified as described previously by Shade and Stopnisek^50^. Briefly, the Bray-Curtis dissimilarity between metagenomic assemblies was estimated at the genus level in a stepwise manner by ranking the genera by detection frequency and average relative abundance (if detection frequency was the same for more than one genus). Proportional Bray-Curtis dissimilarity (fraction of the total average Bray-Curtis dissimilarity) was used to assess the contribution of the ranked genera to the total beta-diversity within these communities. Ecologically relevant core genera were identified by setting a threshold at the point where the addition of further genera contributes to less than 2% of the overall Bray-Curtis dissimilarity.

### Comparative analysis between UBA4765 and Phreatobacter MAGs

16S rRNA gene sequences from the non-redundant (ANI < 99%) UBA4765 MAGs were extracted using barrnap v0.9^51^ (n=20) while 16S rRNA gene sequences from the genus *Phreatobacter* (order: Rhizobiales) were obtained from SILVA v138.1 (n=50) (**Supplemental Table S2**). Pairwise sequence comparisons between 16S rRNA gene sequences were performed using blast 2.5.0^52^ using default alignment parameters. MAGs from genus UBA4765 were compared with *Phreatobacter* genomes from GTDB R207_v2 database and a *Phreatobacter oligotrophus*^53^ genome that was obtained as a part of the DWGC. Pairwise average amino acid identity (AAI) was calculated between UBA4765 and Phreatobacter MAGs using EzAAI v1.2.2^54^ using default parameters. Pairwise ANI between MAGs was calculated using FastANI v1.33^37^ with default parameters.

### Genome annotation and metabolic modeling for UBA4765_DW1549

Gapseq v1.2^55^ was used to construct the metabolic model for select UBA4765 species (UBA4765_DW1549) using the species-level MAG obtained after dereplication at 95% ANI. Annotation was performed with the flags -p all and −l all followed by the function “find-transport” to annotate transporters using default parameters. Draft metabolic model was constructed using these annotations using the function “draft”. Gaps in the model were filled using the “fill” function in gapseq using a custom medium identified for the growth of this organism using the function “medium”. All MAGs within this species (n=42) were used for comparative genome analysis and were annotated using dbCAN^56^, MEROPS^57^, KEGG^58^ and a custom database using METABOLIC v4.0^59^ with the flag -p meta. Biosynthetic gene clusters were identified using antiSMASH v7.0.1^60^ using the flags --cb-general --cb-knownclusters --cb-subclusters --asf --pfam2go --smcog-trees -- genefinding-tool prodigal-m. BacArena^61^ simulations were performed with different carbon and nitrogen sources to verify potential for growth on these sources and these results were used to curate metabolic annotation predictions.

### Statistical analyses

All statistical tests to differentiate between groups used Kruskal Wallis rank sum test and pairwise comparisons between groups were performed using Wilcox test using the stats package on R with a significance cut-off of P<0.05.

## Results and discussion

### Proteobacteria and Patescibacteria represent the most commonly detected phyla in drinking water distribution systems

A total of 13,647 bins were obtained of which 3,170 MAGs/isolate genomes were considered good quality (i.e., quality score > 50) (**Supplemental Table S3)**. Dereplication at 99% ANI cutoff resulted in 1581 good quality non-redundant MAGs which were further clustered at 95% ANI to obtain 1141 species-level clusters. A total of 183 species-level clusters had high-quality draft MAGs as representatives, whereas the remaining 958 species-level clusters had “medium-quality draft” MAGs using MIMAG criteria^42^ due to the absence of one or all genes within the rRNA operon. Of the medium-quality draft species-level MAGs, 739 were greater than 90% complete with less than 5% contamination and 837 had 18 or greater number of unique tRNAs (**Figure 1A**) (**Supplemental Table S4**). Challenges with accurate assembly of rRNA operons and their binning into MAGs is likely the primary issue for a large number of “medium-quality draft” MAGs^62^.

**Figure 1:**
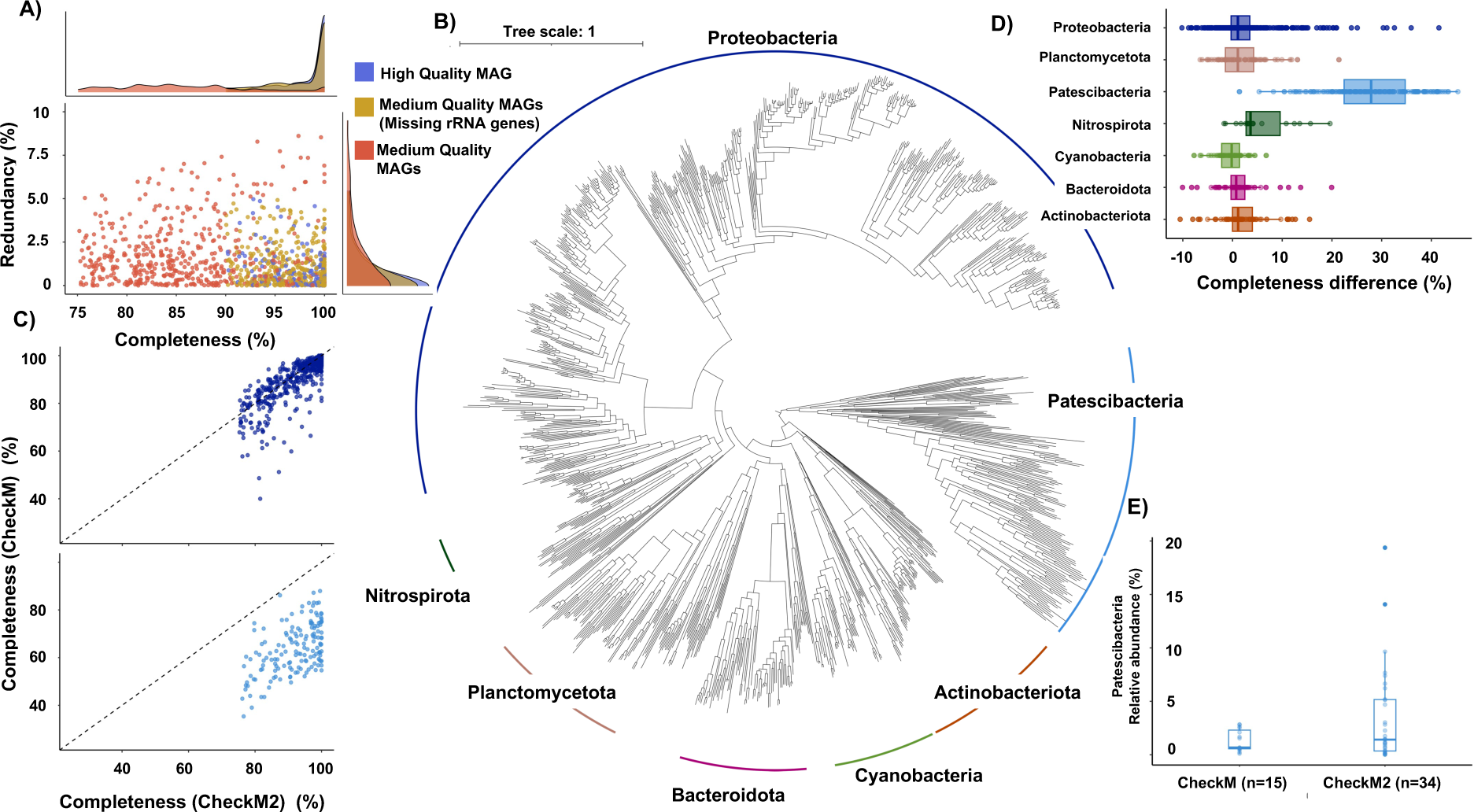
A) Redundancy and Completeness for the 1141 species-level MAGs that form the DWGC. Colors represent the MIMAG quality and density plots depict the redundancy and completeness for the different MIMAG groups. B) Phylogenetic tree of 1141 species-level MAGs capturing the major phyla in drinking water systems. C) Comparisons of CheckM vs CheckM2 estimated completeness for Proteobacteria (top) and Patescibacteria (bottom). D) Difference between CheckM2 and CheckM completeness for the major phyla in the DWGC. E) Relative abundance and number of systems detected for Patescibacteria across different drinking water distribution systems using CheckM Vs CheckM2 while using the same dRep parameters.

Proteobacteria was the most dominant and abundant phylum in the species-level clusters (n = 563, Alphaproteobacteria = 338, Gammaproteobacteria = 225) (**Figure 1B**) (**Supplemental Table S4**) with an average relative abundance of 41.29 ± 27.22% in DWDSs. The 563 proteobacterial species-level clusters contributed to 32.62% of the phylogenetic diversity in the DWGC. Surprisingly, Patescibacteria was the second largest phylum in DWDSs with 156 species-level clusters (**Figure 1B**). Despite the detection of significantly fewer Patescibacteria species relative to Proteobacteria, they capture 24.14% of the phylogenetic diversity. Patescibacteria are seldom detected and described in drinking water studies^14,15^; this could be due to two possible reasons. First, Patescibacteria are underrepresented or missed in gene centric studies (i.e., SSU rRNA gene) due to divergent 16S rRNA gene sequences^63^. Further, Patescibacteria have highly reduced genomes and lack of several ribosomal proteins commonly found in bacteria^64^ and thus genome-centric studies often discard genome bins from this phylum due to lower estimates of genome completeness using CheckM. In contrast, CheckM2 outperforms CheckM in predicting MAG quality for taxa lacking sufficient representation in reference databases and reduced genome sizes while maintaining comparable estimates with CheckM for other taxa^35^(**Figure 1C**). Specifically, the average difference between completeness predictions between CheckM2 and CheckM for Patescibacteria was significantly higher (28.08 ± 8.84%) than for the other prominent phyla in drinking water systems (P<1.6e-10, Pairwise Wilcox test) (**Figure 1D**). Patescibacteria were detected in 42.5% of the systems when competitively mapping reads to MAGs passing quality threshold using CheckM2 estimates compared to detection in 18.75% of metagenomes when relying on CheckM (**Figure 1E**). It is important to note that Patescibacteria had an average relative abundance of 3.73 ± 5.08% which was comparable to other abundant phyla like Actinobacteriota, Planctomycetota and Bacteroidota even if these phyla were observed in more systems than Patescibacteria.

### An uncultured genus of high prevalence was observed in drinking water systems globally

Competitive read mapping from all metagenomes against the 1141 species-level cluster MAGs was used to estimate their detection frequency and relative abundance. The DWGC captures a significantly (*P* = 0.0007) larger proportion of metagenomes from disinfected (56.3 ± 24.89%) as compared to non-disinfected systems (23.5 ± 15.68%) (**Figure 2A**). This result is expected since majority of the metagenomes used for construction of the DWGC were from disinfected (44.7%) as compared to non-disinfected (11.8%) system. Further, approximately 43.5% of metagenomes used in this study were from studies that did not specify the type of disinfectant residual, yet the proportions of reads mapping (54.09 ± 23.83%) to the DWGC was not significantly (*P* = 0.7) different from disinfected systems. It is likely that these metagenomes from studies with no specified disinfection residual could be disinfected systems. The low representativeness of the DWGC for non-disinfected systems suggests that additional effort is required to populate the DWGC with genomes from these systems. It is important to note that non-disinfected systems are significantly more diverse as compared to disinfected systems^3,4,11,13,14^ and thus ensuring a DWGC representative for non-disinfected systems will be challenging.

**Figure 2:**
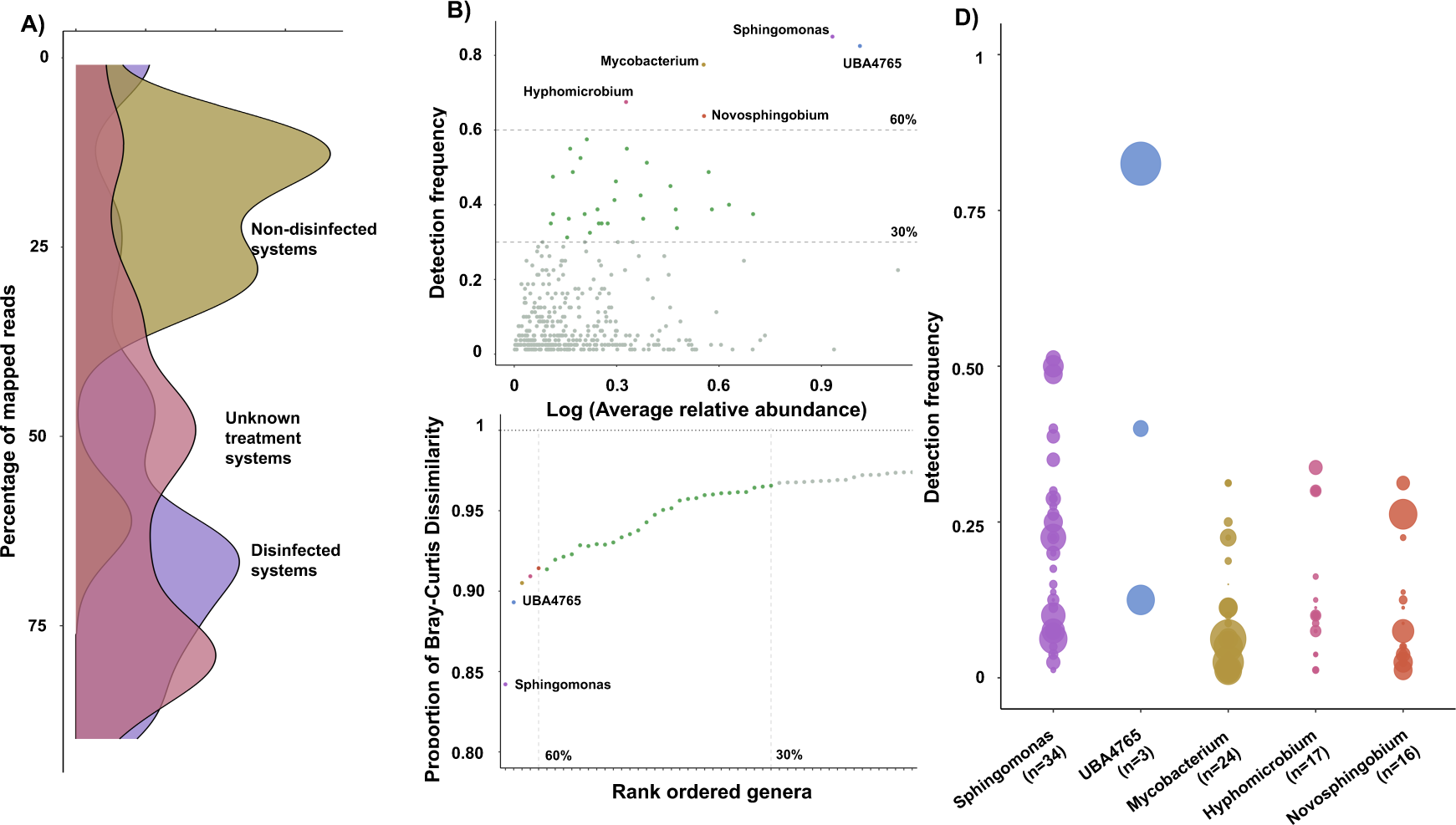
A) Density plot depicting the percentage of mapped reads across all 85 distribution systems against the DWGC separated by treatment strategy. B) Five genera were identified in more than 60% of the systems. B) The five-most frequently detected genera constitute >90% of the proportional Bray Curtis dissimilarity of the entire DWGC. D) Detection frequency of different species within the five genera. Size of the bubble indicates the average relative abundance of these species in the detected systems.

Core microbiome analysis has been widely used to identify and further characterize microbial community members that contribute disproportionately to ecosystem functions^65^. It is important to note that even low abundance taxa can disproportionately contribute to ecosystem function^66^ and their low abundance may be indicative of a unique ecological niche that is consistently observed across multiple DWDSs. We use detection frequency^67^ as opposed to abundance^15^ to identify core taxa since our primary goal was to identify organisms under selection. Furthermore, a recent study also demonstrated that detection frequency-based approach is likely to accurately define core-memberships within an ecosystem^68^. For this analysis, we only used metagenomes where at least 10% of the reads mapped to the DWGC MAGs (n=80). The genera detected in more than 30% and 60% detection frequencies was used to identify “potentially” core genera (**Figure 2B**). A total of 33 genera were detected in more than 30% of the DWDSs of which five were seen in more than 60% (**Figure 2B, Supplementary Table S5**). Consistent with previous findings^10,11,15^, *Sphingomonas* was the most commonly observed genus and was detected in 85% of the DWDSs with an average relative abundance of 7.55 ± 10.9%. Interestingly, an uncultured genus (UBA4765) within the order *Rhizobiales* was just as highly prevalent in the drinking water metagenomes (detection frequency = 82.5%) and at a high average relative abundance (9.29 ± 17.6%). It was first reported by Parks et al. (2017) as a part of multiple uncultured groups with the representative genome for UBA4765 assembled from a drinking water system^69^. A similar trend was observed in the genome distribution data from Liu et al. (2024)^15^ where this genus was observed in 80.2% of their samples.

The 33 genera identified in the above analysis contributed to 96.5% of the dissimilarity between DWDSs with the five genera observed in 60% of the systems contributing to 91.4% of the dissimilarity (**Figure 2B**) estimated as described previously^50^. Interestingly, only two genera (i.e., *Sphingomonas* and UBA4765) explained nearly all of the BC dissimilarity (89.3% of the total dissimilarity) between all DWDSs suggesting a high-level of selection. A total of 34 species-level clusters were identified within the genus *Sphingomona*s, while UBA4765 only had three species-level clusters. Of these UBA4765 species, one species (UBA4765_DW1549) was detected in a large number of systems (82.5%) at a very high relative abundance as well (8.21 ± 16%) (**Figure 2C**); this MAG shares 90.38% ANI with UBA4765 MAG recently reported by Liu et al. (2024)^15^. Considering the global distribution of this MAG in disinfected drinking water systems, it likely represents the genome of a bacterium that is a very important part of the core drinking water microbiome in disinfected DWDSs.

### UBA4765 has been historically misannotated as *Phreatobacter* in 16S rRNA gene sequencing studies

A previously assembled UBA4765 MAG^69^, used as a representative for this genus and family, contains a 16S rRNA gene sequence that matches (100% identity) to that of an organism classified as *Phreatobacter* in the SILVA database (accession number: JQ924015.1.1443). Further, a phreatobacterial sequence from the SILVA database (accession number: JQ684446.1.1463) exhibited 100% sequence identity to the majority of the UBA4765 16S rRNA gene sequences (16 out of 20 sequences extracted from 52 MAGs in this study) across the entire length of the extracted sequence; four of these were partial genes while the remaining were full-length 16S rRNA genes. The same sequence (JQ684446.1.1463) exhibited ∼ 98.5% sequence identity across the entire 16S rRNA gene length against two other 16S rRNA gene sequences extracted from UBA4765 MAGs. It should also be noted that JQ684446.1.1463 was obtained from a DWDS. Pairwise sequence identity comparisons indicated that on average 16S rRNA gene sequences from obtained UBA4765 showed 99.01 ± 1.79% identity with each other (**Figure 3A**). This is expected given that most of these sequences were obtained from UBA4765_DW1549 (19 out of 20). In contrast, a pairwise ANI between sequences classified as *Phreatobacter* in the SILVA database was 91.84 ± 1.79 % and thus are unlikely to be derived from organisms within the same genus^70^. Indeed, we contend that there are currently several sequences placed within the genus “*Phreatobacter*” in the SILVA database that originate from a distinct poorly classified genera (like UBA4765) and leading to mis-annotation of 16S rRNA genes sequenced derived from drinking water systems. While multiple 16S rRNA gene sequencing-based studies have previously detected *Phreatobacter* as one of the most common drinking water microbes^11,71–73^, we only detected it in 5% of metagenomes assembled this study. In contrast, UBA4765 was detected in 82.5% of metagenomes. Therefore, it is highly likely that previous studies reporting “Phreatobacter” in drinking water systems were likely detecting UBA4765. Based on our assessment of the 16S rRNA gene sequence similarities between UBA4765 and the validated species from the genus *Phreatobacter* (ANI values between UBA4765 and Phreatobacter species: *P. oligotrophus* = 93.76 ± 0.74%, *P. stygius* = 93.35 ± 0.75% and *P. cathodhiphilus* = 93.86 ± 0.63%), it appears that UBA4765 and *Phreatobacter* may share the same family (*Phreatobacteraceae*) but are distinct genera^70^ (**Figure 3B**).

**Figure 3:**
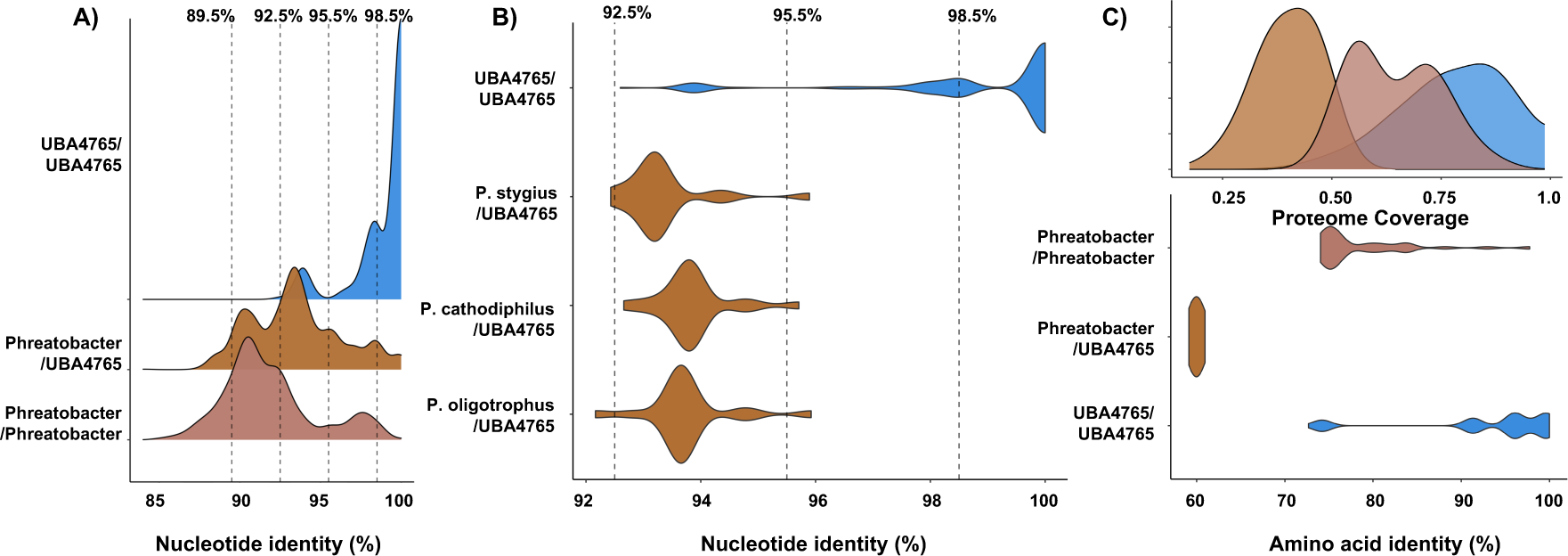
A) Density plot depicting pairwise sequence identity of 16S rRNA gene sequences from UBA4765 MAGs and the sequences of genus *Phreatobacter* obtained from SILVA. Identity cutoffs for species (98.5%), genus (95.5%), family (92.5%) and order (89.5%) are depicted using dotted lines. B) Violin plot depicting identity distribution between 16S rRNA gene sequences from UBA4765 and the three cultured *Phreatobacte*r species (oligotrophus, stygius and cathodiphilus) on LPSN (DSMZ). C) Amino acid identity values from comparing UBA4765 MAGs with genomes from genus *Phreatobacter*. Density plot indicates the proteome coverage between the different groups.

Differentiating between UBA4765 and Phreatobacter is critical because these two genera exhibit significant differences at the genomic level and by extension their functional relevance in drinking water systems. The AAI of 52 MAGs recovered from the genus UBA4765 were compared with nine *Phreatobacter* MAGs obtained from GTDB and this study (**Figure 3C**). *Phreatobacter* MAGs were significantly different from UBA4765. The pairwise AAI values between *Phreatobacter* and UBA4765 (59.98 ± 0.31%) significantly different than the intra-genus values for *Phreatobacter* (78.46 ± 5.52%) and UBA4765 (93.16 ± 8.33%) MAGs (Wilcox pairwise comparison: P<2.2*e-16). Furthermore, the proteome coverage analysis indicated that the gene content of *Phreatobacter* and UBA4765 is very different with a shared proteome of 39.99 ± 6.64% which contrasts significantly with intra-genus shared proteomes for *Phreatobacter* (64.50 ± 9.8%, P<2.2*e-16) and UBA4765 MAGs (77.32 ± 12.12%, P<2.2*e-16).

### Genus UBA4765 consists of discrete populations with varying prevalence and a globally distributed single genomevar

Of the 52 unique MAGs from three different species within the genus UBA4765, 42 belonged to a single species (i.e., UBA4765_DW1549) which were independently assembled and binned from 37 independent DWDSs globally with multiple genomes recovered from some systems likely representing two distinct lineages. While the two lineages exhibit ∼95% ANI with each other, lineage one (UBA4765_DW1549_L1) consists of multiple MAGs that share nearly 99.5% ANI with each other and lineage two (UBA4765_DW1549_L2) consists of multiple MAGs that share ∼98% ANI with each other. Pairwise AAI comparisons between all 52 MAGs resulted in four distinct clusters (**Figure 4A**) representing three distinct species with two lineages within one species; all AAI values shown are relative to comparisons with UBA4765_DW1549_L1. Interestingly, the detection frequency of these species decreases as they become more dissimilar to UBA4765_DW1549_L1 MAGs. The proteome coverage of these groups (species and lineages) also showed variations with more distant clusters exhibiting lower proteome coverage compared to UBA4765_DW1549_L1 MAGs. These differences are not an artifact of MAG completeness, as MAGs with very similar completeness values still display lower proteome coverage.

**Figure 4:**
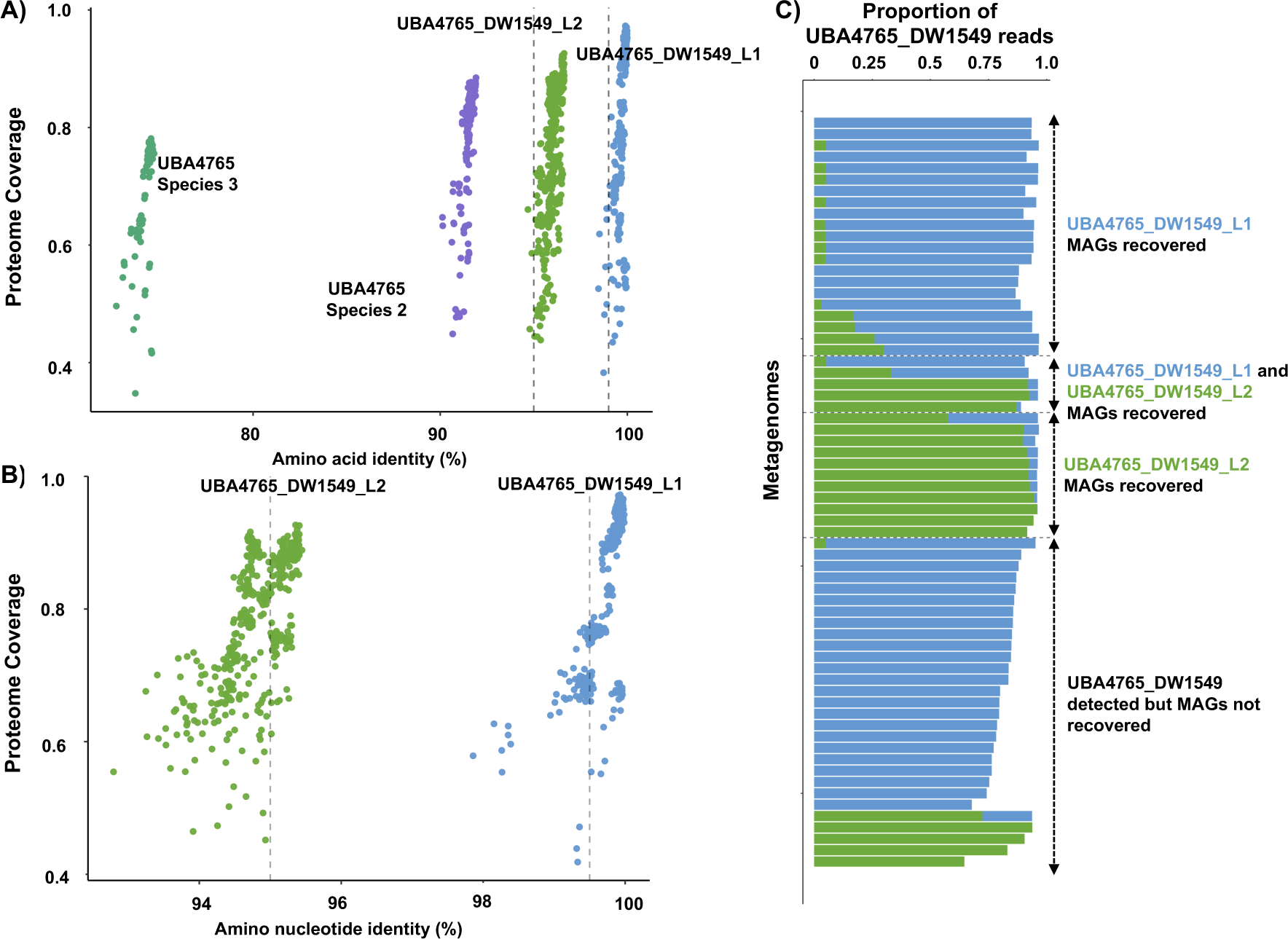
A) Comparisons of the proteome coverage and amino acid identity of genomes within genus UBA4765 with UBA4765 chlorumaquaticus Lineage 1. B) Average nucleotide identity analysis between the two lineages of UBA4765. All comparisons are against UBA4765 Chlorumaquaticus lineage 1. C) Percentage of UBA4765 reads that map to UBA4765_DW1549_L1 and UBA4765_DW1549_L2 MAGS split based on whether UBA4765 MAGs were recovered or detected in metagenomes.

A further evaluation of UBA4765_DW1549 MAGs using ANI analysis (**Figure 4B**) indicated that nearly all MAGs reconstructed within UBA4765_DW1549_L1 likely represent a single genomevar (i.e., all share ANI values greater than 99.5%)^74^. While there are pairwise ANI comparisons outside this threshold for UBA4765_DW1549_L1, this is due to the fragmentation of some of the MAGs used for comparison. In order to evaluate the prevalence of this genomevar across the drinking water metagenomes, we performed competitive mapping of reads mapped to all UBA4765_DW1549 MAGs from metagenomes against all five non-redundant UBA4765_DW1549 MAGs (99% ANI clustering) using a read identify threshold of 99% with a minimum of 75% of the read length mapping. Mapping results indicate that most of these reads mapped to MAGs from UBA4765_DW1549_L1 even if both lineages were detected in DWDS. This suggests that not only is UBA4765_DW1549_L1 being selected for in disinfected drinking water systems, but also that likely there is competitive exclusion between UBA4765_DW1549_L1 and UBA4765_DW1549_L2 as they are never detected at comparable relative abundances in the same DWDS (**Figure 4C**). This genomevar (UBA4765_DW1549_L1) is globally distributed in drinking water metagenomes where it was detected in 60 out of the 66 systems (overall detection frequency: 0.75) where the genus UBA4765 was detected.

It is important to note here that the differentiation of UBA4765_DW1549 into two lineages is provisional. These two lineages share an ANI value right around the 95% threshold with an AAI value above 95% which is why we conclude that they belong to the same species. However, given the trends in differing prevalence of the two lineages within DWDSs, it is also possible that they could represent two different species based on their preference, for as yet unknown, ecosystem conditions and by extension, their phenotypic differences. Therefore, it is likely that they are either: (1) distantly related lineages within the same species and at a point of speciation as indicative of their potential phenotypic differences inferred from competitive mapping or (2) they are two distinct closely related species^75^. Culturing this organism would help us better understand the phenotypic differences between the lineages in order to characterize them better.

### UBA4765_DW1549 genomic content indicates disinfection-mediated selection and metabolism of high relevance to the drinking water ecosystem

Metabolic annotation (**Supplemental Table S6**) suggests that UBA4765_DW1549 is capable of degrading amino acids and utilizing them for growth (**Figure 5A**) with BacArena simulations indicating that it is capable of using 17 of the 20 amino acids which are degraded into the intermediates of the citric acid (TCA) cycle. Interestingly, UBA4765_DW1549 does not possess the metabolic potential to degrade three aromatic amino acids (i.e., phenylalanine, tryptophan and tyrosine) and in general seems to be unable to degrade aromatic compounds. It also has the ability to degrade fatty acids since all the UBA4765_DW1549 MAGs contain the beta-oxidation pathway (KEGG:M00087) responsible for breaking down fatty acids into acetyl-CoA which can enter the TCA cycle. This organism also contains multiple peptidases and carbohydrate hydrolyzing enzymes capable of degrading complex molecules like chitin, xylan, peptidoglycan, and peptides (to sugars and amino acids) which can be utilized for growth. The presence of these pathways and the BacArena simulation results indicate that UBA4765_DW1549 could likely utilize decaying biomass within distribution systems for growth (i.e., nectrotrophic lifestyle)^13,76^. UBA4765_DW1549 has the ability to utilize C1 carbons like formate and CO for metabolism like other members within the order Rhizobiales and their genomes harbor haloacid dehalogenase (KEGG:K01560) required to degrade halogenated compounds (e.g., 2-haloacids)

**Figure 5:**
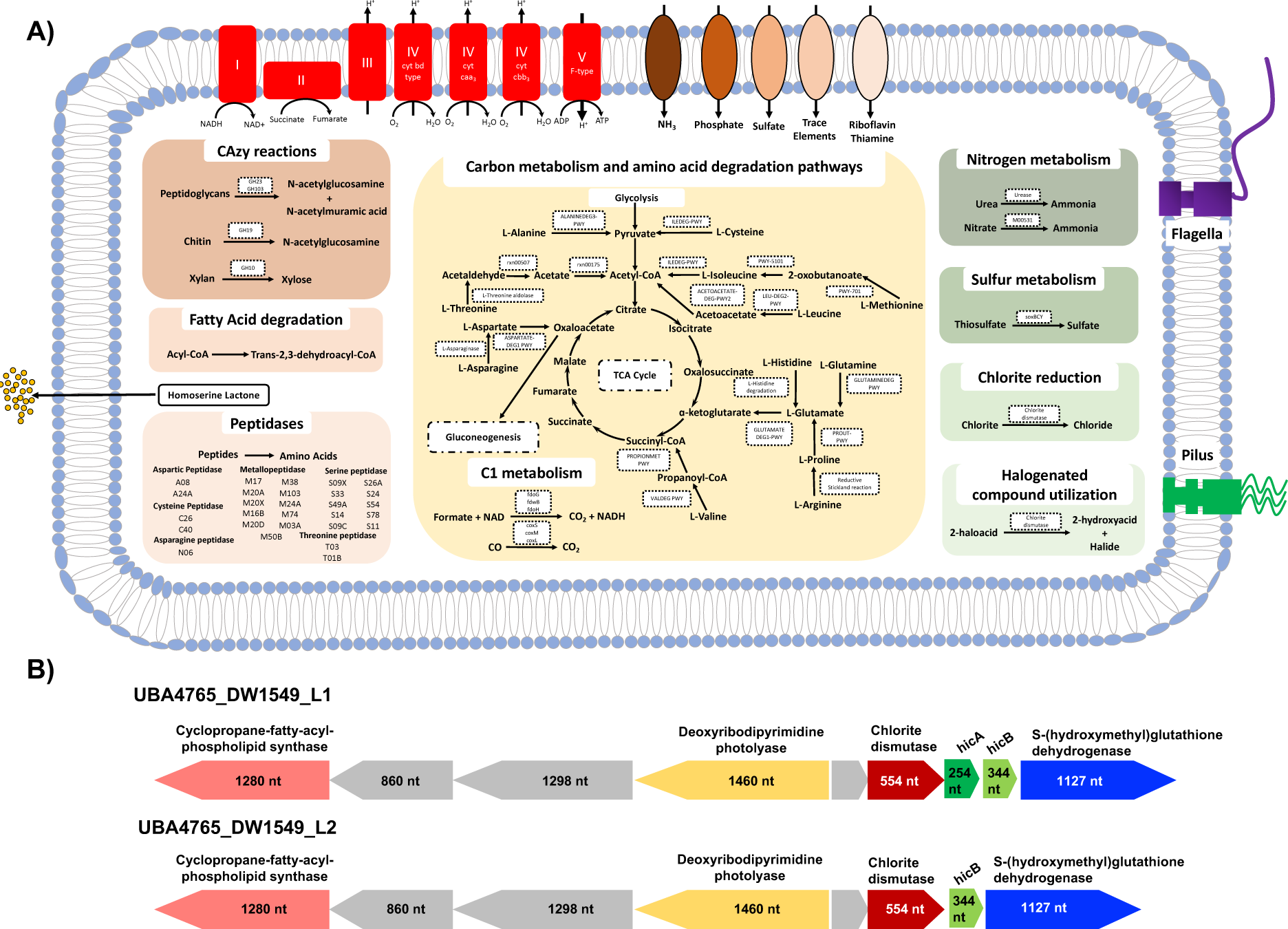
A) Predicted metabolism of UBA4765_DW1549 determined using METABOLIC, Antismash and Gapseq. B) Chlorite dismutase and neighboring genes in UBA4765_DW1549_L1 and UBA4765_DW1549_L2.

UBA4765_DW1549 also exhibits metabolic traits that are of high relevance to the disinfected drinking water environment. It has the metabolic potential to synthesize homoserine lactones which has been associated with quorum sensing and biofilm formation^77^. Based on the Gapseq construction of the metabolic model, this organism is incapable of producing riboflavin and thiamine. Therefore, adaptation to a biofilm environment serves as an opportunity to obtain these essential nutrients via proximity to organisms that produce them while also providing protection from disinfectant residuals. Interestingly, UBA4765_DW1549 MAGs include a gene encoding for chlorite dismutase (KEGG:K09162) which is implicated in the degradation of chlorite. The chlorite dismutase gene was detected in vast majority of independently assembled UBA4765_DW1549 MAGs in both lineages without the perchlorate reductase (PCRA) gene; this could suggest selection since the occurrence of the gene has been linked to chlorite presence in the environment^78^. Chlorite dismutase gene is only observed in 1% of the genomes and 5% of the genera in the NCBI taxonomy and is not widely distributed^78^. Inspection of the neighborhood of the chlorite dismutase genes further highlighted genetic potential that may allow for persistence in a disinfected DWDS and fine-scale differences between the two lineages that may explain the selection of UBA4765_DW1549_L1 over UBA4765_DW1549_L2. Nearly all MAGs from both lineages encoded S-(hydroxymethyl)glutathione dehydrogenase (KEGG:K00121) which is associated with formaldehyde detoxification but also with redox regulation^79^ and could play a role in oxidative stress response. Further, all UBA4765_DW1549 MAGs encode a deoxyribodipyrimidine photolyase (KEGG: K01669) which is associated with UV radiation-induced DNA damage^80^. Similarly, all UBA4765_DW1549 encode cyclopropane-fatty-acyl-phospholipid synthase (KEGG: K00574) responsible for the synthesis of cyclopropane fatty acids (CFA) which is associated bacterial membrane protection against environmental stressors^81^ and CFAs have also been detected in DWDS^82^.

Nearly all UBA4765_DW1549_L1 MAGs genes encode for toxin-antitoxin system HicAB (**Figure 5B**) immediately downstream of the chlorite dismutase gene. The HicAB toxin-antioxin system is associated with persister/dormancy phenotypes allowing the cell to function under high stress conditions^83^. Interestingly, UBA4765_DW1549_L2 only had the hicB gene downstream of chlorite dismutase, with the hicA gene found on a different contig; this was not due to contig fragmentation. Thus, it is plausible that the HicAB toxin-antioxin system is more tightly regulated in UBA4765_DW1549_L1 as compared to UBA4765_DW1549_L2. In addition to the potential differential regulation of persister phenotype, the ability to oxidize thiosulfate was only detected in in UBA4765_DW1549_L1 and not in UBA4765_DW1549_L2. The ability to cycle sulfur compounds would be particularly advantageous in a chlorine stressed environment^84^. These differences need to be further studied to better understand their role in fitness of both two lineages considering UBA4765_DW1549_L1 is far more prevalent. It is interesting that all of these genes associated with stress tolerance, DNA repair, persister phenotype are co-located with the chlorite dismutase gene. This could suggest disinfection-mediated selection for UBA4765_DW1549 in disinfected DWDSs. Indeed, we observed significant increase in the relative abundance of UBA4765_DW1549 post-disinfection in multiple datasets^85–91^.

### Proposal of a new name for UBA4765_DW1549

We demonstrate that UBA4765_DW1549 has likely been persistently detected in most culture independent investigations of DWDSs but was incorrectly annotated as *Phreatobacter*. This species represents the only uncharacterized group of organisms that was detected in vast majority of drinking water metagenomes (i.e., >80%) and at very high relative abundance, suggesting that it likely constitutes a vast majority of the microbial community (and possibly biomass) in disinfected DWDSs. Further, it is remarkable that even within this select group, there are signs of selection. Specifically, a single genomevar within this species is globally distributed and harbors traits that indicate disinfection-mediated selection (i.e., chlorite dismutase without PCRA) along with co-localized genes that confer additional advantages in a stressed environment. This along with the ability to utilize decaying biomass and the ability to form biofilms makes the detailed physiological characterization of UBA4765_DW1549 critical for understanding microbial growth and biofilm formation in DWDSs. Indeed, if cultured, UBA4765_DW1549 would represent the ideal model organism for understanding the ecology and physiology of the drinking water microbiome in disinfected systems.

To facilitate systematic future investigations of this important bacterium, we propose a provisional name for the uncultured genus UBA4765 as “*Raskinella*” and for the species UBA4765_DW1549 as “*Raskinella chlorumaquaticus*” (Pending approval from SeqCode^92^). “*Raskinella*” is named after Dr. Lutgarde Raskin for her extensive contributions to the field of drinking water microbiology and microbial ecology. The species name *Chlorumaquaticus* (Chlorum.aqua.ti.cus) is attributed to the observation that this bacterium is only detected in disinfected drinking water systems and appears to be selected for through the process of drinking water disinfection.

## Data availability

The non-redundant MAGs that form the basis of the DWGC and species-level representative MAGs are available on FigShare (DOI: https://doi.org/10.6084/m9.figshare.c.7245403.v1)

## Acknowledgements

This research was supported by the National Science Foundation (NSF) (CBET Award Number: 2220792).

## References

1) Hull, N. M.; Ling, F.; Pinto, A. J.; Albertsen, M.; Jang, H. G.; Hong, P.-Y.; Konstantinidis, K. T.; LeChevallier, M.; Colwell, R. R.; Liu, W.-T. Drinking Water Microbiome Project: Is It Time? Trends in Microbiology 2019, 27 (8), 670–677. 10.1016/j.tim.2019.03.011.

2) Proctor, C. R.; Hammes, F. Drinking Water Microbiology—from Measurement to Management. Current Opinion in Biotechnology 2015, 33, 87–94. 10.1016/j.copbio.2014.12.014.

3) Gabrielli, M.; Dai, Z.; Delafont, V.; Timmers, P. H. A.; van der Wielen, P. W. J. J.; Antonelli, M.; Pinto, A. J. Identifying Eukaryotes and Factors Influencing Their Biogeography in Drinking Water Metagenomes. Environ. Sci. Technol. 2023, 57 (9), 3645–3660. 10.1021/acs.est.2c09010.

4) Hegarty, B.; Dai, Z.; Raskin, L.; Pinto, A.; Wigginton, K.; Duhaime, M. A Snapshot of the Global Drinking Water Virome: Diversity and Metabolic Potential Vary with Residual Disinfectant Use. Water Research 2022, 218, 118484. 10.1016/j.watres.2022.118484.

5) Ke, Y.; Sun, W.; Jing, Z.; Zhao, Z.; Xie, S. Seasonal Variations of Microbial Community and Antibiotic Resistome in a Suburb Drinking Water Distribution System in a Northern Chinese City. Journal of Environmental Sciences 2023, 127, 714–725. 10.1016/j.jes.2022.07.001.

6) Pinto, A. J.; Schroeder, J.; Lunn, M.; Sloan, W.; Raskin, L. Spatial-Temporal Survey and Occupancy-Abundance Modeling To Predict Bacterial Community Dynamics in the Drinking Water Microbiome. mBio 2014, 5 (3), 10.1128/mbio.01135-14. 10.1128/mbio.01135-14.

7) Pinto, A. J.; Xi, C.; Raskin, L. Bacterial Community Structure in the Drinking Water Microbiome Is Governed by Filtration Processes. Environ. Sci. Technol. 2012, 46 (16), 8851–8859. 10.1021/es302042t.

8) Prest, E. I.; Hammes, F.; van Loosdrecht, M. C. M.; Vrouwenvelder, J. S. Biological Stability of Drinking Water: Controlling Factors, Methods, and Challenges. Frontiers in Microbiology 2016, 7.

9) Proctor, C.; Garner, E.; Hamilton, K. A.; Ashbolt, N. J.; Caverly, L. J.; Falkinham, J. O.; Haas, C. N.; Prevost, M.; Prevots, D. R.; Pruden, A.; Raskin, L.; Stout, J.; Haig, S.-J. Tenets of a Holistic Approach to Drinking Water-Associated Pathogen Research, Management, and Communication. Water Research 2022, 211, 117997. 10.1016/j.watres.2021.117997.

10) Santos, Q. M. B. los; Schroeder, J. L.; Sevillano-Rivera, M. C.; Sungthong, R.; Ijaz, U. Z.; Sloan, W. T.; Pinto, A. J. Emerging Investigators Series: Microbial Communities in Full-Scale Drinking Water Distribution Systems – a Meta-Analysis. Environ. Sci.: Water Res. Technol. 2016, 2 (4), 631–644. 10.1039/C6EW00030D.

11) Thom, C.; Smith, C. J.; Moore, G.; Weir, P.; Ijaz, U. Z. Microbiomes in Drinking Water Treatment and Distribution: A Meta-Analysis from Source to Tap. Water Research 2022, 212, 118106. 10.1016/j.watres.2022.118106.

12) Berry, D.; Xi, C.; Raskin, L. Microbial Ecology of Drinking Water Distribution Systems. Current Opinion in Biotechnology 2006, 17 (3), 297–302. 10.1016/j.copbio.2006.05.007.

13) Dai, Z.; Sevillano-Rivera, M. C.; Calus, S. T.; Bautista-de los Santos, Q. M.; Eren, A. M.; van der Wielen, P. W. J. J.; Ijaz, U. Z.; Pinto, A. J. Disinfection Exhibits Systematic Impacts on the Drinking Water Microbiome. Microbiome 2020, 8 (1), 42. 10.1186/s40168-020-00813-0.

14) Gomez-Alvarez, V.; Siponen, S.; Kauppinen, A.; Hokajärvi, A.-M.; Tiwari, A.; Sarekoski, A.; Miettinen, I. T.; Torvinen, E.; Pitkänen, T. A Comparative Analysis Employing a Gene- and Genome-Centric Metagenomic Approach Reveals Changes in Composition, Function, and Activity in Waterworks with Different Treatment Processes and Source Water in Finland. Water Research 2023, 229, 119495. 10.1016/j.watres.2022.119495.

15) Liu, H.; Jiao, P.; Guan, L.; Wang, C.; Zhang, X.-X.; Ma, L. Functional Traits and Health Implications of the Global Household Drinking-Water Microbiome Retrieved Using an Integrative Genome-Centric Approach. Water Research 2024, 250, 121094. 10.1016/j.watres.2023.121094.

16) Potgieter, S. C.; Dai, Z.; Venter, S. N.; Sigudu, M.; Pinto, A. J. Microbial Nitrogen Metabolism in Chloraminated Drinking Water Reservoirs. mSphere 2020, 5 (2), 10.1128/msphere.00274-20. 10.1128/msphere.00274-20.

17) Sevillano, M.; Dai, Z.; Calus, S.; Bautista-de los Santos, Q. M.; Eren, A. M.; van der Wielen, P. W. J. J.; Ijaz, U. Z.; Pinto, A. J. Differential Prevalence and Host-Association of Antimicrobial Resistance Traits in Disinfected and Non-Disinfected Drinking Water Systems. Science of The Total Environment 2020, 749, 141451. 10.1016/j.scitotenv.2020.141451.

18) Tiwari, A.; Gomez-Alvarez, V.; Siponen, S.; Sarekoski, A.; Hokajärvi, A.-M.; Kauppinen, A.; Torvinen, E.; Miettinen, I. T.; Pitkänen, T. Bacterial Genes Encoding Resistance Against Antibiotics and Metals in Well-Maintained Drinking Water Distribution Systems in Finland. Front. Microbiol. 2022, 12. 10.3389/fmicb.2021.803094.

19) Huang, D.; Yuan, M. M.; Chen, J.; Zheng, X.; Wong, D.; Alvarez, P. J. J.; Yu, P. The Association of Prokaryotic Antiviral Systems and Symbiotic Phage Communities in Drinking Water Microbiomes. ISME Communications 2023, 3 (1), 46. 10.1038/s43705-023-00249-1.

19) Solize, V. Genome Centric and Flow Cytometric Characterization of the Boston Water Microbiome. Ph.D. Dissertation, >Northeastern University, Boston, MA 2022.

21) Chen, S.; Zhou, Y.; Chen, Y.; Gu, J. Fastp: An Ultra-Fast All-in-One FASTQ Preprocessor. Bioinformatics 2018, 34 (17), i884–i890. 10.1093/bioinformatics/bty560.

22) Li, H. Aligning Sequence Reads, Clone Sequences and Assembly Contigs with BWA-MEM. arXiv May 26, 2013. 10.48550/arXiv.1303.3997.

23) Vasimuddin, Md.; Misra, S.; Li, H.; Aluru, S. Efficient Architecture-Aware Acceleration of BWA-MEM for Multicore Systems. In 2019 IEEE International Parallel and Distributed Processing Symposium (IPDPS); 2019; pp 314–324. 10.1109/IPDPS.2019.00041.

24) Danecek, P.; Bonfield, J. K.; Liddle, J.; Marshall, J.; Ohan, V.; Pollard, M. O.; Whitwham, A.; Keane, T.; McCarthy, S. A.; Davies, R. M.; Li, H. Twelve years of SAMtools and BCFtools. GigaScience 2021, 10 (2), giab008. 10.1093/gigascience/giab008.

25) Quinlan, A. R.; Hall, I. M. BEDTools: A Flexible Suite of Utilities for Comparing Genomic Features. Bioinformatics 2010, 26 (6), 841–842. 10.1093/bioinformatics/btq033.

26) Nurk, S.; Meleshko, D.; Korobeynikov, A.; Pevzner, P. A. metaSPAdes: A New Versatile Metagenomic Assembler. Genome Res. 2017, 27 (5), 824–834. 10.1101/gr.213959.116.

27) Langmead, B.; Salzberg, S. L. Fast Gapped-Read Alignment with Bowtie 2. Nat Methods 2012, 9 (4), 357–359. 10.1038/nmeth.1923.

28) Kang, D. D.; Li, F.; Kirton, E.; Thomas, A.; Egan, R.; An, H.; Wang, Z. MetaBAT 2: An Adaptive Binning Algorithm for Robust and Efficient Genome Reconstruction from Metagenome Assemblies. PeerJ 2019, 7, e7359. 10.7717/peerj.7359.

29) Alneberg, J.; Bjarnason, B. S.; de Bruijn, I.; Schirmer, M.; Quick, J.; Ijaz, U. Z.; Lahti, L.; Loman, N. J.; Andersson, A. F.; Quince, C. Binning Metagenomic Contigs by Coverage and Composition. Nat Methods 2014, 11 (11), 1144–1146. 10.1038/nmeth.3103.

30) Nissen, J. N.; Johansen, J.; Allesøe, R. L.; Sønderby, C. K.; Armenteros, J. J. A.; Grønbech, C. H.; Jensen, L. J.; Nielsen, H. B.; Petersen, T. N.; Winther, O.; Rasmussen, S. Improved Metagenome Binning and Assembly Using Deep Variational Autoencoders. Nat Biotechnol 2021, 39 (5), 555–560. 10.1038/s41587-020-00777-4.

31) Eren, A. M.; Kiefl, E.; Shaiber, A.; Veseli, I.; Miller, S. E.; Schechter, M. S.; Fink, I.; Pan, J. N.; Yousef, M.; Fogarty, E. C.; Trigodet, F.; Watson, A. R.; Esen, Ö. C.; Moore, R. M.; Clayssen, Q.; Lee, M. D.; Kivenson, V.; Graham, E. D.; Merrill, B. D.; Karkman, A.; Blankenberg, D.; Eppley, J. M.; Sjödin, A.; Scott, J. J.; Vázquez-Campos, X.; McKay, L. J.; McDaniel, E. A.; Stevens, S. L. R.; Anderson, R. E.; Fuessel, J.; Fernandez-Guerra, A.; Maignien, L.; Delmont, T. O.; Willis, A. D. Community-Led, Integrated, Reproducible Multi-Omics with Anvi’o. Nat Microbiol 2021, 6 (1), 3–6. 10.1038/s41564-020-00834-3.

32) Sevillano, M.; Vosloo, S.; Cotto, I.; Dai, Z.; Jiang, T.; Santiago Santana, J. M.; Padilla, I. Y.; Rosario-Pabon, Z.; Velez Vega, C.; Cordero, J. F.; Alshawabkeh, A.; Gu, A.; Pinto, A. J. Spatial-Temporal Targeted and Non-Targeted Surveys to Assess Microbiological Composition of Drinking Water in Puerto Rico Following Hurricane Maria. Water Research X 2021, 13, 100123. 10.1016/j.wroa.2021.100123.

33) Vosloo, S.; Huo, L.; Anderson, C. L.; Dai, Z.; Sevillano, M.; Pinto, A. Evaluating de Novo Assembly and Binning Strategies for Time Series Drinking Water Metagenomes. Microbiology Spectrum 2021, 9 (3), e01434-21. 10.1128/Spectrum.01434-21.

34) Parks, D. H.; Imelfort, M.; Skennerton, C. T.; Hugenholtz, P.; Tyson, G. W. CheckM: Assessing the Quality of Microbial Genomes Recovered from Isolates, Single Cells, and Metagenomes. Genome Res. 2015, 25 (7), 1043–1055. 10.1101/gr.186072.114.

35) Chklovski, A.; Parks, D. H.; Woodcroft, B. J.; Tyson, G. W. CheckM2: A Rapid, Scalable and Accurate Tool for Assessing Microbial Genome Quality Using Machine Learning. Nat Methods 2023, 20 (8), 1203–1212. 10.1038/s41592-023-01940-w.

36) Olm, M. R.; Brown, C. T.; Brooks, B.; Banfield, J. F. dRep: A Tool for Fast and Accurate Genomic Comparisons That Enables Improved Genome Recovery from Metagenomes through de-Replication. The ISME Journal 2017, 11 (12), 2864–2868. 10.1038/ismej.2017.126.

37) Jain, C.; Rodriguez-R, L. M.; Phillippy, A. M.; Konstantinidis, K. T.; Aluru, S. High Throughput ANI Analysis of 90K Prokaryotic Genomes Reveals Clear Species Boundaries. Nat Commun 2018, 9 (1), 5114. 10.1038/s41467-018-07641-9.

38) Almeida, A.; Nayfach, S.; Boland, M.; Strozzi, F.; Beracochea, M.; Shi, Z. J.; Pollard, K. S.; Sakharova, E.; Parks, D. H.; Hugenholtz, P.; Segata, N.; Kyrpides, N. C.; Finn, R. D. A Unified Catalog of 204,938 Reference Genomes from the Human Gut Microbiome. Nat Biotechnol 2021, 39 (1), 105–114. 10.1038/s41587-020-0603-3.

39) Chaumeil, P.-A.; Mussig, A. J.; Hugenholtz, P.; Parks, D. H. GTDB-Tk v2: Memory Friendly Classification with the Genome Taxonomy Database. Bioinformatics 2022, 38 (23), 5315–5316. 10.1093/bioinformatics/btac672.

40) Schwengers, O.; Jelonek, L.; Dieckmann, M. A.; Beyvers, S.; Blom, J.; Goesmann, A. Bakta: Rapid and Standardized Annotation of Bacterial Genomes via Alignment-Free Sequence Identification. Microbial Genomics 2021, 7 (11), 000685. 10.1099/mgen.0.000685.

41) Seemann, T. Prokka: Rapid Prokaryotic Genome Annotation. Bioinformatics 2014, 30 (14), 2068–2069. 10.1093/bioinformatics/btu153.

42) Bowers, R. M.; Kyrpides, N. C.; Stepanauskas, R.; Harmon-Smith, M.; Doud, D.; Reddy, T. B. K.; Schulz, F.; Jarett, J.; Rivers, A. R.; Eloe-Fadrosh, E. A.; Tringe, S. G.; Ivanova, N. N.; Copeland, A.; Clum, A.; Becraft, E. D.; Malmstrom, R. R.; Birren, B.; Podar, M.; Bork, P.; Weinstock, G. M.; Garrity, G. M.; Dodsworth, J. A.; Yooseph, S.; Sutton, G.; Glöckner, F. O.; Gilbert, J. A.; Nelson, W. C.; Hallam, S. J.; Jungbluth, S. P.; Ettema, T. J. G.; Tighe, S.; Konstantinidis, K. T.; Liu, W.-T.; Baker, B. J.; Rattei, T.; Eisen, J. A.; Hedlund, B.; McMahon, K. D.; Fierer, N.; Knight, R.; Finn, R.; Cochrane, G.; Karsch-Mizrachi, I.; Tyson, G. W.; Rinke, C.; Lapidus, A.; Meyer, F.; Yilmaz, P.; Parks, D. H.; Murat Eren, A.; Schriml, L.; Banfield, J. F.; Hugenholtz, P.; Woyke, T. Minimum Information about a Single Amplified Genome (MISAG) and a Metagenome-Assembled Genome (MIMAG) of Bacteria and Archaea. Nat Biotechnol 2017, 35 (8), 725–731. 10.1038/nbt.3893.

43) Capella-Gutiérrez, S.; Silla-Martínez, J. M.; Gabaldón, T. trimAl: A Tool for Automated Alignment Trimming in Large-Scale Phylogenetic Analyses. Bioinformatics 2009, 25 (15), 1972–1973. 10.1093/bioinformatics/btp348.

44) Stamatakis, A. RAxML Version 8: A Tool for Phylogenetic Analysis and Post-Analysis of Large Phylogenies. Bioinformatics 2014, 30 (9), 1312–1313. 10.1093/bioinformatics/btu033.

45) Letunic, I.; Bork, P. Interactive Tree of Life (iTOL) v6: Recent Updates to the Phylogenetic Tree Display and Annotation Tool. Nucleic Acids Research 2024, gkae268. 10.1093/nar/gkae268.

46) Faith, D. P. Conservation Evaluation and Phylogenetic Diversity. Biological Conservation 1992, 61 (1), 1–10. 10.1016/0006-3207(92)91201-3.

47) Bittinger K (2020). _abdiv: Alpha and Beta Diversity Measures. R package version 0.2.0. https://CRAN.R-project.org/package=abdiv.

48) R Core Team (2023). R: A Language and Environment for Statistical Computing. R Foundation for Statistical Computing, Vienna, Austria. https://www.R-project.org/.

49) Aroney, S. T. N., Newell, R. J. P., Nissen, J., Camargo, A. P., Tyson, G. W., & Woodcroft, B. J. (2024). CoverM: Read coverage calculator for metagenomics (v0.7.0). Zenodo. 10.5281/zenodo.10531254

50) Shade, A.; Stopnisek, N. Abundance-Occupancy Distributions to Prioritize Plant Core Microbiome Membership. Current Opinion in Microbiology 2019, 49, 50–58. 10.1016/j.mib.2019.09.008

51) Seemann T barrnap 0.9 : rapid ribosomal RNA prediction https://github.com/tseemann/barrnap

52) Altschul, S. F.; Gish, W.; Miller, W.; Myers, E. W.; Lipman, D. J. Basic Local Alignment Search Tool. Journal of Molecular Biology 1990, 215 (3), 403–410. 10.1016/S0022-2836(05)80360-2.

53) Tóth, E. M.; Vengring, A.; Homonnay, Z. G.; Kéki, Zs.; Spröer, C.; Borsodi, A. K.; Márialigeti, K.; Schumann, P. Phreatobacter Oligotrophus Gen. Nov., Sp. Nov., an Alphaproteobacterium Isolated from Ultrapure Water of the Water Purification System of a Power Plant. International Journal of Systematic and Evolutionary Microbiology 2014, 64 (Pt_3), 839–845. 10.1099/ijs.0.053843-0.

54) Kim, D.; Park, S.; Chun, J. Introducing EzAAI: A Pipeline for High Throughput Calculations of Prokaryotic Average Amino Acid Identity. J Microbiol. 2021, 59 (5), 476–480. 10.1007/s12275-021-1154-0.

55) Zimmermann, J.; Kaleta, C.; Waschina, S. Gapseq: Informed Prediction of Bacterial Metabolic Pathways and Reconstruction of Accurate Metabolic Models. Genome Biology 2021, 22 (1), 81. 10.1186/s13059-021-02295-1.

56) Yin, Y.; Mao, X.; Yang, J.; Chen, X.; Mao, F.; Xu, Y. dbCAN: A Web Resource for Automated Carbohydrate-Active Enzyme Annotation. Nucleic Acids Research 2012, 40 (W1), W445–W451. 10.1093/nar/gks479.

57) Rawlings, N. D.; Barrett, A. J.; Thomas, P. D.; Huang, X.; Bateman, A.; Finn, R. D. The MEROPS Database of Proteolytic Enzymes, Their Substrates and Inhibitors in 2017 and a Comparison with Peptidases in the PANTHER Database. Nucleic Acids Research 2018, 46 (D1), D624–D632. 10.1093/nar/gkx1134.

58) Kanehisa, M.; Furumichi, M.; Sato, Y.; Kawashima, M.; Ishiguro-Watanabe, M. KEGG for Taxonomy-Based Analysis of Pathways and Genomes. Nucleic Acids Research 2023, 51 (D1), D587–D592. 10.1093/nar/gkac963.

59) Zhou, Z.; Tran, P. Q.; Breister, A. M.; Liu, Y.; Kieft, K.; Cowley, E. S.; Karaoz, U.; Anantharaman, K. METABOLIC: High-Throughput Profiling of Microbial Genomes for Functional Traits, Metabolism, Biogeochemistry, and Community-Scale Functional Networks. Microbiome 2022, 10 (1), 33. 10.1186/s40168-021-01213-8.

60) Blin, K.; Shaw, S.; Augustijn, H. E.; Reitz, Z. L.; Biermann, F.; Alanjary, M.; Fetter, A.; Terlouw, B. R.; Metcalf, W. W.; Helfrich, E. J. N.; van Wezel, G. P.; Medema, M. H.; Weber, T. antiSMASH 7.0: New and Improved Predictions for Detection, Regulation, Chemical Structures and Visualisation. Nucleic Acids Research 2023, 51 (W1), W46–W50. 10.1093/nar/gkad344.

61) Bauer, E.; Zimmermann, J.; Baldini, F.; Thiele, I.; Kaleta, C. BacArena: Individual-Based Metabolic Modeling of Heterogeneous Microbes in Complex Communities. PLOS Computational Biology 2017, 13 (5), e1005544. 10.1371/journal.pcbi.1005544.

62) Zhang, Y.; Ji, P.; Wang, J.; Zhao, F. RiboFR-Seq: A Novel Approach to Linking 16S rRNA Amplicon Profiles to Metagenomes. Nucleic Acids Research 2016, 44 (10), e99. 10.1093/nar/gkw165.

63) Brown, C. T.; Hug, L. A.; Thomas, B. C.; Sharon, I.; Castelle, C. J.; Singh, A.; Wilkins, M. J.; Wrighton, K. C.; Williams, K. H.; Banfield, J. F. Unusual Biology across a Group Comprising More than 15% of Domain Bacteria. Nature 2015, 523 (7559), 208–211. 10.1038/nature14486.

64) Castelle, C. J.; Banfield, J. F. Major New Microbial Groups Expand Diversity and Alter Our Understanding of the Tree of Life. Cell 2018, 172 (6), 1181–1197. 10.1016/j.cell.2018.02.016.

65) Shade, A.; Handelsman, J. Beyond the Venn Diagram: The Hunt for a Core Microbiome. Environmental Microbiology 2012, 14 (1), 4–12. 10.1111/j.1462-2920.2011.02585.x.

66) Jousset, A.; Bienhold, C.; Chatzinotas, A.; Gallien, L.; Gobet, A.; Kurm, V.; Küsel, K.; Rillig, M. C.; Rivett, D. W.; Salles, J. F.; van der Heijden, M. G. A.; Youssef, N. H.; Zhang, X.; Wei, Z.; Hol, W. H. G. Where Less May Be More: How the Rare Biosphere Pulls Ecosystems Strings. ISME J 2017, 11 (4), 853–862. 10.1038/ismej.2016.174.

67) Neu, A. T.; Allen, E. E.; Roy, K. Defining and Quantifying the Core Microbiome: Challenges and Prospects. Proceedings of the National Academy of Sciences 2021, 118 (51), e2104429118. 10.1073/pnas.2104429118.

68) Custer, G. F.; Gans, M.; van Diepen, L. T. A.; Dini-Andreote, F.; Buerkle, C. A. Comparative Analysis of Core Microbiome Assignments: Implications for Ecological Synthesis. mSystems 2023, 8 (1), e01066-22. 10.1128/msystems.01066-22.

69) Parks, D. H.; Rinke, C.; Chuvochina, M.; Chaumeil, P.-A.; Woodcroft, B. J.; Evans, P. N.; Hugenholtz, P.; Tyson, G. W. Recovery of Nearly 8,000 Metagenome-Assembled Genomes Substantially Expands the Tree of Life. Nat Microbiol 2017, 2 (11), 1533–1542. 10.1038/s41564-017-0012-7.

70) Konstantinidis, K. T.; Rosselló-Móra, R.; Amann, R. Uncultivated Microbes in Need of Their Own Taxonomy. ISME J 2017, 11 (11), 2399–2406. 10.1038/ismej.2017.113.

71) Perrin, Y.; Bouchon, D.; Delafont, V.; Moulin, L.; Héchard, Y. Microbiome of Drinking Water: A Full-Scale Spatio-Temporal Study to Monitor Water Quality in the Paris Distribution System. Water Research 2019, 149, 375–385. 10.1016/j.watres.2018.11.013.

72) A. Chavarria, K.; I. Gonzalez, C.; Goodridge, A.; Saltonstall, K.; L. Nelson, K. Bacterial Communities in a Neotropical Full-Scale Drinking Water System Including Intermittent Piped Water Supply, from Sources to Taps. Environmental Science: Water Research & Technology 2023. 10.1039/D3EW00224A.

73) Vosloo, S.; Huo, L.; Chauhan, U.; Cotto, I.; Gincley, B.; Vilardi, K. J.; Yoon, B.; Bian, K.; Gabrielli, M.; Pieper, K. J.; Stubbins, A.; Pinto, A. J. Gradual Recovery of Building Plumbing-Associated Microbial Communities after Extended Periods of Altered Water Demand during the COVID-19 Pandemic. Environ. Sci. Technol. 2023, 57 (8), 3248–3259. 10.1021/acs.est.2c07333.

74) Rodriguez-R, L. M.; Conrad, R. E.; Viver, T.; Feistel, D. J.; Lindner, B. G.; Venter, S. N.; Orellana, L. H.; Amann, R.; Rossello-Mora, R.; Konstantinidis, K. T. An ANI Gap within Bacterial Species That Advances the Definitions of Intra-Species Units. mBio 2023, 15 (1), e02696-23. 10.1128/mbio.02696-23.

75) Zhao, C.; Shi, Z. J.; Pollard, K. S. Pitfalls of Genotyping Microbial Communities with Rapidly Growing Genome Collections. cels 2023, 14 (2), 160-176.e3. 10.1016/j.cels.2022.12.007.

76) Chatzigiannidou, I.; Props, R.; Boon, N. Drinking Water Bacterial Communities Exhibit Specific and Selective Necrotrophic Growth. npj Clean Water 2018, 1 (1), 1–4. 10.1038/s41545-018-0023-9.

77) Parsek, M. R.; Greenberg, E. P. Acyl-Homoserine Lactone Quorum Sensing in Gram-Negative Bacteria: A Signaling Mechanism Involved in Associations with Higher Organisms. Proceedings of the National Academy of Sciences 2000, 97 (16), 8789–8793. 10.1073/pnas.97.16.8789.

78) Barnum, T. P.; Coates, J. D. Chlorine Redox Chemistry Is Widespread in Microbiology. ISME J 2022, 1–14. 10.1038/s41396-022-01317-5.

79) Imber, M.; Pietrzyk-Brzezinska, A. J.; Antelmann, H. Redox Regulation by Reversible Protein *S*-Thiolation in Gram-Positive Bacteria. Redox Biology 2019, 20, 130–145. 10.1016/j.redox.2018.08.017.

80) Matallana-Surget, S.; Douki, T.; Cavicchioli, R.; Joux, F. Remarkable Resistance to UVB of the Marine Bacterium Photobacterium Angustum Explained by an Unexpected Role of Photolyase. Photochem Photobiol Sci 2009, 8 (9), 1313–1320. 10.1039/b902715g.

81) Zhu, X.; Guo, Z.; Wang, N.; Liu, J.; Zuo, Y.; Li, K.; Song, C.; Song, Y.; Gong, C.; Xu, X.; Yuan, F.; Zhang, L. Environmental Stress Stimulates Microbial Activities as Indicated by Cyclopropane Fatty Acid Enhancement. Science of The Total Environment 2023, 873, 162338. 10.1016/j.scitotenv.2023.162338.

82) Smith, C. A.; Phiefer, C. B.; Macnaughton, S. J.; Peacock, A.; Burkhalter, R. S.; Kirkegaard, R.; White, D. C. Quantitative Lipid Biomarker Detection of Unculturable Microbes and Chlorine Exposure in Water Distribution System Biofilms. Water Research 2000, 34 (10), 2683–2688. 10.1016/S0043-1354(00)00028-2.

83) Harms, A.; Brodersen, D. E.; Mitarai, N.; Gerdes, K. Toxins, Targets, and Triggers: An Overview of Toxin-Antitoxin Biology. Molecular Cell 2018, 70 (5), 768–784. 10.1016/j.molcel.2018.01.003.

84) Gray, M. J.; Wholey, W.-Y.; Jakob, U. Bacterial Responses to Reactive Chlorine Species. Annual Review of Microbiology 2013, 67 (volume 67, 2013), 141–160. 10.1146/annurev-micro-102912-142520.

85) Dias, M. F.; da Rocha Fernandes, G.; Cristina de Paiva, M.; Christina de Matos Salim, A.; Santos, A. B.; Amaral Nascimento, A. M. Exploring the Resistome, Virulome and Microbiome of Drinking Water in Environmental and Clinical Settings. Water Research 2020, 174, 115630. 10.1016/j.watres.2020.115630.

86) Gu, Q.; Sun, M.; Lin, T.; Zhang, Y.; Wei, X.; Wu, S.; Zhang, S.; Pang, R.; Wang, J.; Ding, Y.; Liu, Z.; Chen, L.; Chen, W.; Lin, X.; Zhang, J.; Chen, M.; Xue, L.; Wu, Q. Characteristics of Antibiotic Resistance Genes and Antibiotic-Resistant Bacteria in Full-Scale Drinking Water Treatment System Using Metagenomics and Culturing. Frontiers in Microbiology 2022, 12.

87) Shi, P.; Jia, S.; Zhang, X.-X.; Zhang, T.; Cheng, S.; Li, A. Metagenomic Insights into Chlorination Effects on Microbial Antibiotic Resistance in Drinking Water. Water Research 2013, 47 (1), 111–120. 10.1016/j.watres.2012.09.046.

88) Jia, S.; Shi, P.; Hu, Q.; Li, B.; Zhang, T.; Zhang, X.-X. Bacterial Community Shift Drives Antibiotic Resistance Promotion during Drinking Water Chlorination. Environ. Sci. Technol. 2015, 49 (20), 12271–12279. 10.1021/acs.est.5b03521.

89) Chao, Y.; Ma, L.; Yang, Y.; Ju, F.; Zhang, X.-X.; Wu, W.-M.; Zhang, T. Metagenomic Analysis Reveals Significant Changes of Microbial Compositions and Protective Functions during Drinking Water Treatment. Sci Rep 2013, 3 (1), 3550. 10.1038/srep03550.

90) Zhao, Q.; He, H.; Gao, K.; Li, T.; Dong, B. Fate, Mobility, and Pathogenicity of Drinking Water Treatment Plant Resistomes Deciphered by Metagenomic Assembly and Network Analyses. Science of The Total Environment 2022, 804, 150095. 10.1016/j.scitotenv.2021.150095.

91) Ke, Y.; Sun, W.; Jing, Z.; Zhao, Z.; Xie, S. Seasonal Variations of Microbial Community and Antibiotic Resistome in a Suburb Drinking Water Distribution System in a Northern Chinese City. Journal of Environmental Sciences 2023, 127, 714–725. 10.1016/j.jes.2022.07.001.

92) Hedlund, B. P.; Chuvochina, M.; Hugenholtz, P.; Konstantinidis, K. T.; Murray, A. E.; Palmer, M.; Parks, D. H.; Probst, A. J.; Reysenbach, A.-L.; Rodriguez-R, L. M.; Rossello-Mora, R.; Sutcliffe, I. C.; Venter, S. N.; Whitman, W. B. SeqCode: A Nomenclatural Code for Prokaryotes Described from Sequence Data. Nat Microbiol 2022, 7 (10), 1702–1708. 10.1038/s41564-022-01214-9.

